# Gene-drive-mediated extinction is thwarted by evolution of sib mating

**DOI:** 10.1101/558924

**Authors:** James J Bull, Christopher H Remien, Stephen M Krone

## Abstract

Genetic engineering combined with CRISPR technology has developed to the point that gene drives can, in theory, be engineered to cause extinction in countless species. Success of extinction programs now rests on the possibility of resistance evolution, which is largely unknown. For CRISPR technology, resistance may take many forms, from mutations in the nuclease target sequence to specific types of non-random population structures that limit the drive. We develop mathematical models of various deviations from random mating to consider escapes from extinction-causing gene drives. We use a version of Maynard Smith’s haystack model to show that population structure can enable drive-free subpopulations to be maintained against gene drives. Our main emphasis, however, is sib mating in the face of recessive-lethal and Y-chromosome drives. Sib mating easily evolves in response to both kinds of gene drives and maintains mean fitness above 0, with equilibrium fitness depending on the level of inbreeding depression. Environmental determination of sib mating (as might stem from population density crashes) can also maintain mean fitness above 0. Translation of mean fitness into population size depends on ecological details, so understanding mean fitness evolution and dynamics is merely the first step in predicting extinction. Nonetheless, these results point to possible escapes from gene drive-mediated extinctions that lie beyond the control of genome engineering.

## 1 Introduction

Engineered gene drives offer an exciting new technology for the possible control of pests and vector-borne diseases, and which might even be used to rescue wildlife species from the edge of extinction. The selective advantage of some drives is so powerful that they can be used to cause species extinction, but many potential applications propose using them more benignly, to deliver a harmless genetic cargo throughout a species.

It seems paradoxical that natural selection can favor genes that cause extinction, but the theory indicating such possibilities is over half a century old (Prout, 1953; Bruck, 1957; Lewontin, 1958; Hamilton, 1967). Engineering to implement these systems remained the challenge for decades, but the insight of Burt (2003) combined with CRISPR technology has led to a revolution in interest (Sinkins and Gould, 2006; Gould, 2008; Burt, 2014; Esvelt et al., 2014; Unckless et al., 2015); laboratory experiments have now shown the feasibility of various implementations (Akbari et al., 2013; Gantz and Bier, 2015; Kyrou et al., 2018).

The pace at which engineering methods have enabled gene drive construction has vastly exceeded our experience with implementations, so that we stand poised to introduce gene drives on a massive scale without appreciating how they might fail or deviate from expectations. Given the demonstrated success of engineered gene drives in experimental populations, the most obvious basis of possible failure now becomes the evolution of resistance, the focus of this paper. At a minimum, resistance would limit coverage of the population by a gene drive; at worst, resistance would fully reverse a drive’s effect. Furthermore, the evolution of resistance to one implementation may thwart subsequent implementations, so early failures may have long-term ramifications for later interventions. There is thus an imperative to understand resistance evolution before implementing gene drives on a wide scale.

Resistance may take many forms. Some forms may be specific to the mechanistic underpinnings of the gene drive implementation, others may operate largely independent of the drive mechanism. For a homing endonuclease gene, an obvious form of resistance is mutation in the target sequence recognized by the nuclease (Burt, 2003). Resistance that, at a molecular level, blocks gene drive expression or interferes with its operation will be difficult to predict or study except empirically, in the context of specific applications. Other types of resistance, especially those that transcend mechanistic details of the drive, may be more amenable to *a priori* analysis.

Here we specifically consider population structure as a foundation for resistance – how specific types of non-random mating will work and evolve to thwart extinction-causing gene drives, which entail the strongest selection for resistance. We develop two classes of models. The first is a metapopulation model with simplified dynamics regulating local extinctions (from a gene drive), recolonization, and interactions between drive and non-drive populations. It is merely a modified version of Maynard Smith’s original haystack model (Maynard Smith, 1964). The second class allows sib mating in response to a gene drive introduction. We compare the effects of sib mating and its evolution for both Y-chromosome gene drive, which does not kill individuals, and for recessive-lethal gene drive, which does kill. Sib mating evolves in response to both types of gene drive, partially or completely blocking the drive, but the outcomes differ quantitatively between the two types of drive. We also consider non-evolutionary forms of sib mating that may thwart a drive through effects on population structure.

## 2 Consequences of extreme population structure: the haystack model revisited

Many intended uses of gene drives rely on the drive suppressing or even extinguishing populations. The populations most suited to this end are unstructured with random-mating. As is well known from the decades-old theoretical literature on group selection of cooperative (altruistic) traits, a structured population is protected against selfish elements, and the degree of protection depending on quantitative details of fitness effects, migration rates, and group extinction rates (Williams, 1966; Coyne et al., 1997; Leigh, 2010). When the selfish element is a lethal gene drive, it accelerates extinction of the subpopulations in which it resides, but the then-empty patches are recolonized disproportionately by the altruists lacking the selfish element.

To illustrate this process in a highly simplified but intuitively tractable form, we modify the original haystack model of Maynard Smith (Maynard Smith, 1964), with parallels to Hamilton and May’s dispersal model (Hamilton and May, 1977). We imagine many small populations, each inhabiting patches (islands) in a large habitat of many patch sites; i.e., a metapopulation (Fig. 1). Migrants from one population can colonize other sites. There are two types of populations: those consisting purely of wild-type individuals (B, for beneficial) and those with at least some gene-drive individuals (S, for selfish). In the spirit of the haystack model, gene drive spread within a local population is considered so rapid that any patch with even a few gene drive individuals is immediately converted to type S. This separation of time scales is an essential feature of metapopulation models and allows for a consolidation of state variables. Although migrants from B cannot convert S populations and thus can be ignored, the reverse migration (rate *β*_*SB*_) is highly effective because of the gene drive effect. Furthermore, high values of *β*_*SB*_ represent a near absence of population structure. The model does not specifically include or even require genetics, the key within-patch process being the rapid takeover of B patches by S individuals when they invade B.

**Figure 1:**
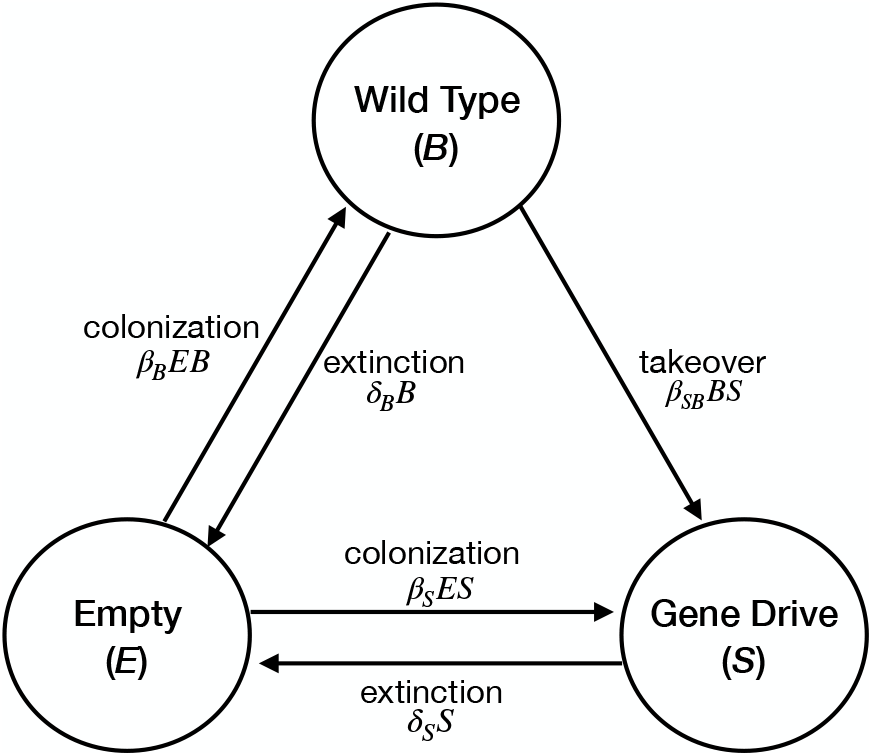
A haystack model of population structure. Three types of patches exist: empty, selfish (containing individuals with the gene drive), and wild-type or beneficial. The rates at which one patch type is converted to another are given by the terms on the arrows, corresponding to equations (1).

**Figure 2:**
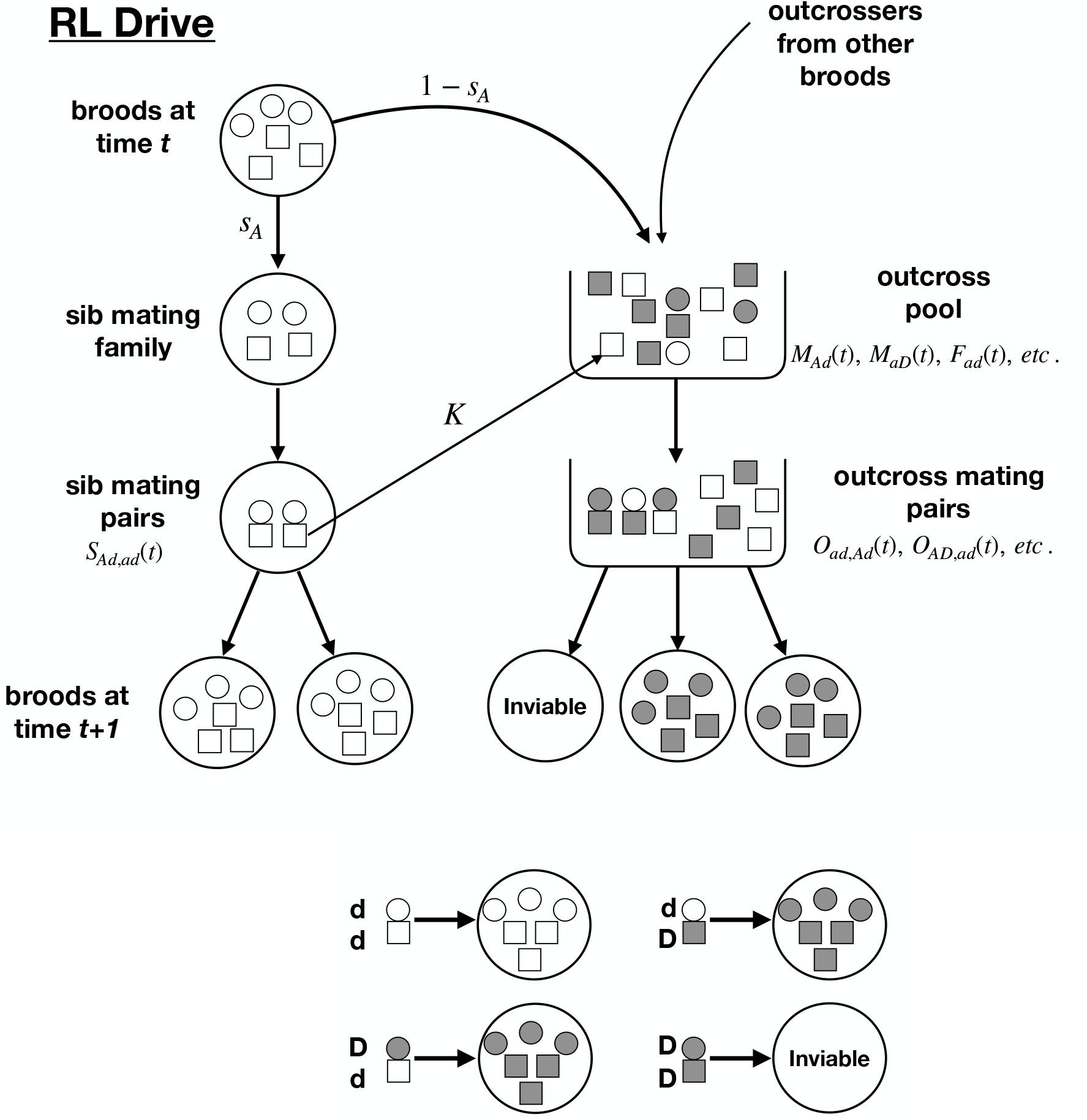
Schematic of the recessive lethal drive model with haploid individuals that enjoy a brief diploid phase for mating. Females are circles, males are squares. Gray indicates the drive allele (D), clear indicates the non-drive allele (d). Top: Shown in the pedigree of large circles on the left is the life cycle of a brood that resulted from a mating of an *d* female and an *d* male. The female sib mating allele (not depicted) determines the fraction of the brood (here *s*_*A*_) that is sib mated and the fraction 1 - *s*_*A*_ that go to the outcross pool along with gametes from other broods that are available to outcross. A fraction *K* of males that sib mate a sister also join the outcross pool to possibly mate some more. Bottom: The mating of a non-drive female and a non-drive male results in all offspring lacking the drive; the mating of a drive female and a drive male results in no offspring; a mating between one drive parent and one non-drive parent results in all offspring carrying the drive. The influence of *σ* is not shown.

Differential equations describing the dynamics are presented in equations (1), with variables and parameters defined in table 1.

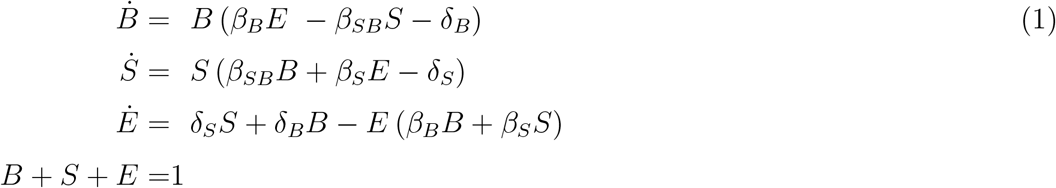

**Table 1:** Model variables and parameters

| Notation | Description |
| --- | --- |
| Variables |  |
| $B$ | frequency of patches with pure wild-type individuals |
| $S$ | frequency of patches with at least some gene drive individuals |
| $E$ | frequency of empty patches |
| Parameters |  |
| $\delta_S$ | extinction rate of S patches |
| $\delta_B$ | extinction rate of B patches |
| $\beta_B$ | rate parameter for B patches colonizing E patches |
| $\beta_S$ | rate parameter for S patches colonizing E patches |
| $\beta_{SB}$ | rate parameter for S patches invading and converting B patches |

Notes: all variables are confined to $[0,1]$ . All parameter values must be positive.

### Parameter constraints

Biology dictates that our interest is confined to parameter values where B alone can persist but S alone cannot (resulting in the conditions *β*_*B*_ *> δ*_*B*_, *δ*_*S*_ *> β*_*S*_), a higher colonization rate of empty patches by B than by S (*β*_*B*_ *> β*_*S*_), and a higher extinction rate of S patches than of B patches (*δ*_*S*_ *> δ*_*B*_). Biology also dictates that all parameters must be non-negative, but there are no upper limits except as imposed by the aforementioned constraints. With these conditions, the system has two relevant equilibria: an equilibrium in which S is absent.

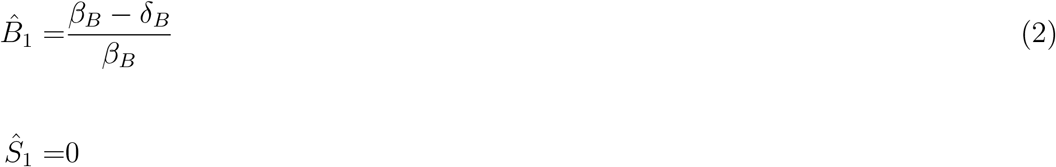

and an internal equilibrium of

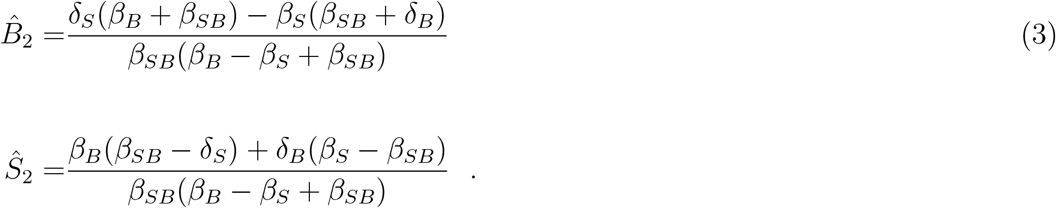

Stability conditions imply that 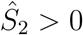 is required for S to invade. Systematically testing which values satisfy (3) across the range of (0.01, 2.02) for all 5 parameters (and when the parameter constraints are met), approximately 15% of the parameter space allows B and S coexistence; for the other 85%, S simply cannot invade. S cannot drive B extinct with these deterministic processes because there is always a threshold value of S below which B is so unaffected by S (by low colonization) that it persists. However, a choice of *β*_*SB*_ sufficiently high can push the equilibrium value of *B* to such a low value that B would not persist in any habitat with a finite number of patches.

Although the haystack model was originally used for insight about group selection, our version here is probably better thought of in the usual metapopulation sense as a model of implicit spatial structure. The resulting spatial segregation is enough to allow coexistence of *B* and *S* (under appropriate conditions on the parameters). What fraction of natural populations satisfy the structural requisites to contain a gene drive by local extinction remains to be seen. However, North et al. (2013) simulated an extinction-causing gene drive in a spatial population with many details specific to mosquito biology. They found that small patches of mosquitoes could escape the extinction wave, suggestive of the patch model here.

## 3 Sib mating

Groups *per se* aren’t the only structures that may operate to thwart an extinguishing gene drive. Structure may exist at the family level in the form of inbreeding. A previous theoretical study found that selfing can be selected in response to a recessive lethal gene drive and that selfing limits the potential for extinction of the population targeted by the drive (Bull, 2016). This result is worrying for gene drive implementations, but there were two hopeful outcomes from that work. First, although evolution of even partial selfing could prevent gene drive fixation, mean fitness of the targeted population was limited by the magnitude of inbreeding depression. Thus, mean fitness remained low if inbreeding depression was high, preserving much of the intended effect of the gene drive. Second, selfing could sometimes only evolve by major mutations, not small ones. That study considered only recessive lethal drive, and the latter result was evaluated only for the case of drive in one sex only. We thus expand upon that work here to address two questions for a different form of inbreeding: (i) Does sib mating also evolve as a block to extinguishing gene drives, and can it evolve in small steps? (ii) Do recessive lethal drives and Y-drives equally favor sib mating?

All models assume a life cycle with sexual haploids: male and female parents mate and produce a brief diploid phase, which then undergoes meiosis to produce haploid offspring. Gene drive operates in the diploid phase (regardless of which parent contributes the drive allele in the recessive-lethal model). Drive is always complete, with 100% of the progeny receiving the drive allele. The locus controlling sib mating has 2 alleles and is unlinked to the drive locus.

### Genotypes and phenotypes

For models with genetic control of sib mating, the family’s level of sib mating is controlled by the mother’s genotype at the *A/a* locus (Table 2). Sib mating is limited to families that produce both sexes; with Y-drive, some families are all sons with no sisters to mate. In all trials illustrated here, sib mating in allele ‘*a*’ was set to 0 – purely outcrossing – and ‘*A*’ provided some sib mating.

**Table 2:** Maternal genotype control of sib mating rate

| Maternal genotype | Proportion daughters mated by brothers |
| --- | --- |
| $a$ | $s_a (=0)$ |
| $A$ | $s_A$ |

A non-genetic (‘ecological’) class of models tested here allows the level of sib mating to change dynamically with mean fitness 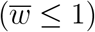. The biological justification is that, as mean fitness declines, so will population density, and siblings may increasingly provide the only potential mates. Here, there is no genetic variation for sib mating (all genotypes are ‘a’ or, equivalently, *s*_*a*_ = *s*_*A*_), and we used an exponential sib-mating function with a single parameter (*c*) to control steepness:

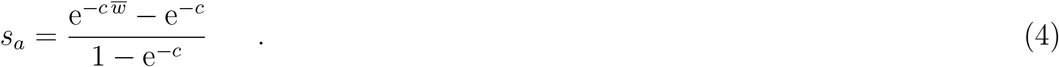

### Drive genetics

Two models of drive are studied. Recessive-lethal drive (alleles *D/d*) operates so that if either (but only one) parent carries *D*, all offspring inherit *D*. If both parents carry *D*, no offspring are produced. That is, *D* is a recessive lethal only in the brief diploid phase. For the model of Y drive, all females carry X, but males carry either Y or Z chromosome. Of a male that carries Y, half his offspring inherit the Y and half inherit the mother’s X. Of a male that carries Z, all his offspring inherit the Z and are thus sons in a brood with no sisters, and hence no possibility of sib mating.

### Inbreeding depression

The models assume that the relative brood size of parents who are sibs is *σ*, typically lower than that of outcrossed offspring (*σ <* 1). (Inbreeding depression would be represented as the decrement in fitness, *δ* = 1 −*σ*.) Inbreeding depression is assumed to be invariant throughout the evolutionary process. In real systems, inbreeding depression is often partially purged upon extended inbreeding, but allowing inbreeding depression to be static is a reasonable starting point and, if anything, provides a conservative measure of the vulnerability of gene drive systems to be suppressed by inbreeding.

### Male reproductive versatility

The net reproductive output of sons from a family with sib mating can be modeled in different ways, each of which may be observed in nature. At one extreme, sons who mate their sisters may then go on to join the random mating pool with no adverse consequence to their abilities in the outcrossing pool. At the other extreme, sons who mate their sisters are forever lost to the outcrossing pool, as if there is a brief time in which all mating occurs and an individual can be at only one place during that time. This latter process, of sons being ‘discounted,’ is conveniently represented by assuming the extreme case that the fraction of a family’s sons lost to the outcrossing pool is the same as the fraction of daughters mated by sons. We use *K* to represent the probability that a sister-mating male also participates in the random-mating pool.

### Four models

The preceding account has identified two fundamental biological differences that require specific models:

a. The allele with drive is an autosomal recessive lethal or a Y (Z) chromosome
b. Sib mating is controlled genetically or ecologically.

Addressing these variables in all combinations, there are four models to study. Equations are given in the Appendix; all models consist of difference equations that assume discrete generations. Drive was assumed to be complete in all cases.

### Recessive lethal drive

#### Genetic control of sib mating

There are minimally 16 viable family types that must be counted: 4 initiated from sib mating, 12 from outcrossing (Appendix; formally, there are 20 mating types, but 4 produce no progeny). The drive allele (D) is present in 8 of the outcrossed family types, and for these families, all progeny carry D so any sib mating is non-productive. Sib-mating rates depend on the mother’s genotype at the *a/A* locus, and the rate of sib mating induced by ‘*A*’ was varied systematically in different trials. Both the drive allele D and sib-mating allele A were introduced at low frequency at the beginning of each trial. Sib mating rates were unaffected by the presence of D in the brood, so D led to offspring death from sib mating.

##### Equilibrium

We describe equilibrium outcomes based on mean fitness, which in our case is the average number of daughters per mother within a generation – all daughters are assumed to be mated. (For the equations in the Appendix, this calculation is 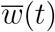 when *b* = 2.) These numerical studies revealed that the sib mating allele always evolved, even when it effected only a small level of sib mating (e.g., *s*_*A*_ = 0.01). However, average fitness at equilibrium was strongly dependent on the magnitude of sib mating encoded by allele ‘A’ up to a value of *s*_*A*_ = 0.5 (fig. 3). With *s*_*A*_ *<* 0.5, allele ‘A’ fixed and mean fitness remained below *σ*. If instead, *s*_*A*_ *>* 0.5, allele ‘A’ remained polymorphic, and mean fitness equalled *σ*. In all cases, mean fitness was bounded by *σ*, the fraction of maximum brood size attained with parents who were sibs.)

**Figure 3:**
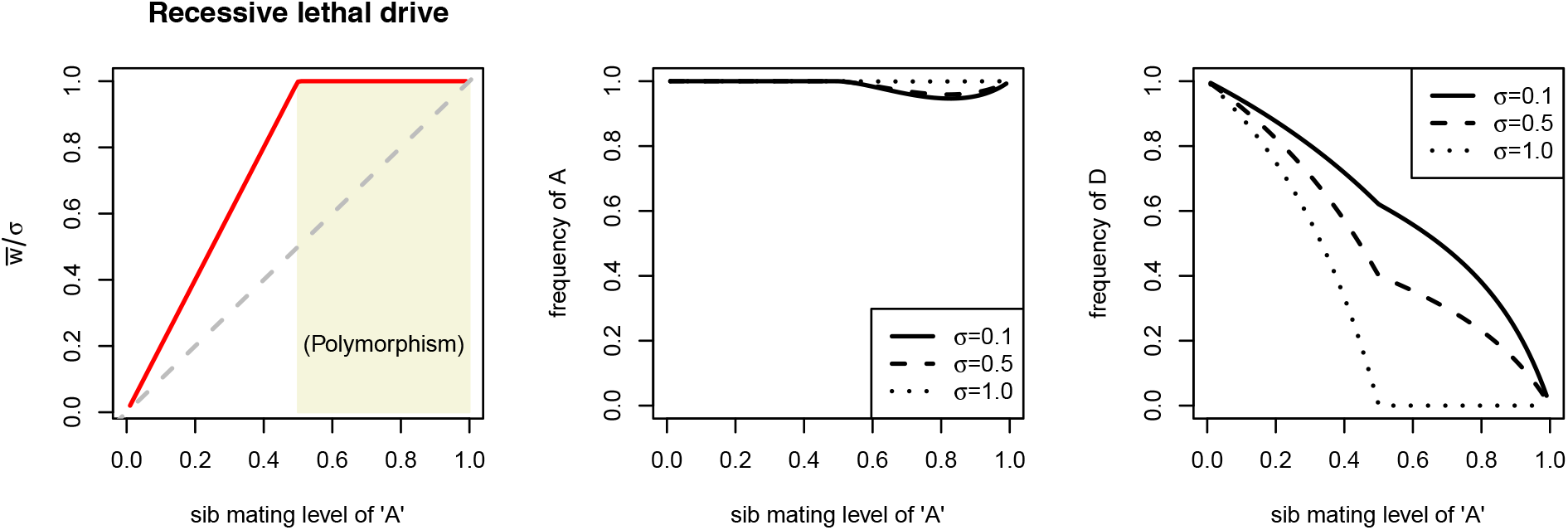
Equilibrium properties for the genetic control of sib mating with recessive lethal drive. The horizontal axis in all panels is *s*_*A*_, the probability of sib-mating for allele ‘A’. All three panels are for the same runs, merely illustrating different properties. (Left) Relative mean fitness attained across different *s*_*A*_ values. 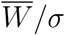 is mean fitness scaled by the fraction of maximum brood size attained by parents who were sibs. That this ratio was never observed to exceed one means that *σ* sets the upper limit on mean fitness. The shaded area shows the region in which the sib-mating allele (A) remained polymorphic. The dashed gray line is an isocline at which the y-value equals the x-value. (Middle) Equilibrium frequency of allele ‘A’. The allele invariably fixes for *s*_*A*_ values up to 0.5, but remains polymorphic at higher *s*_*A*_ values. The value of *σ* has little effect on the frequency. (Right) Equilibrium frequency of the drive allele, D, for different *s*_*A*_ values. Except at the extremes of no sib mating and complete sib mating, the final frequency of D is strongly affected by both *s*_*A*_ and *σ*. These runs assumed full male discounting (*K* = 0).

The foregoing results apply to complete male discounting (*K* = 0 – males who mate their sisters are lost to the random pool). In the absence of male discounting, mean fitness was observed to exceed *σ* when *s*_*A*_ *>* 0.5, but the largest mean fitness observed was 1.3*σ*, and the effect diminished as *σ* increased above 0.5. In comparison to the case of complete male discounting, a higher mean fitness with no male discounting is understandable because of the male fitness gained when sib-mating males later join the random mating pool.

The main qualitative result is that, although sib mating evolves in response to a lethal gene drive, mean fitness is approximately bounded by the fitness consequences of sib mating (*σ*) and also somewhat bounded by the magnitude of sib mating allowed by the genetics.

#### Ecological adjustment of sib mating

In this model, all evolution is limited to the drive locus because genetic variation in sib mating is not required when the level of sib mating increases ‘ecologically.’ We assumed that sib mating increases as mean fitness declines (fig. 4, left).

**Figure 4:**
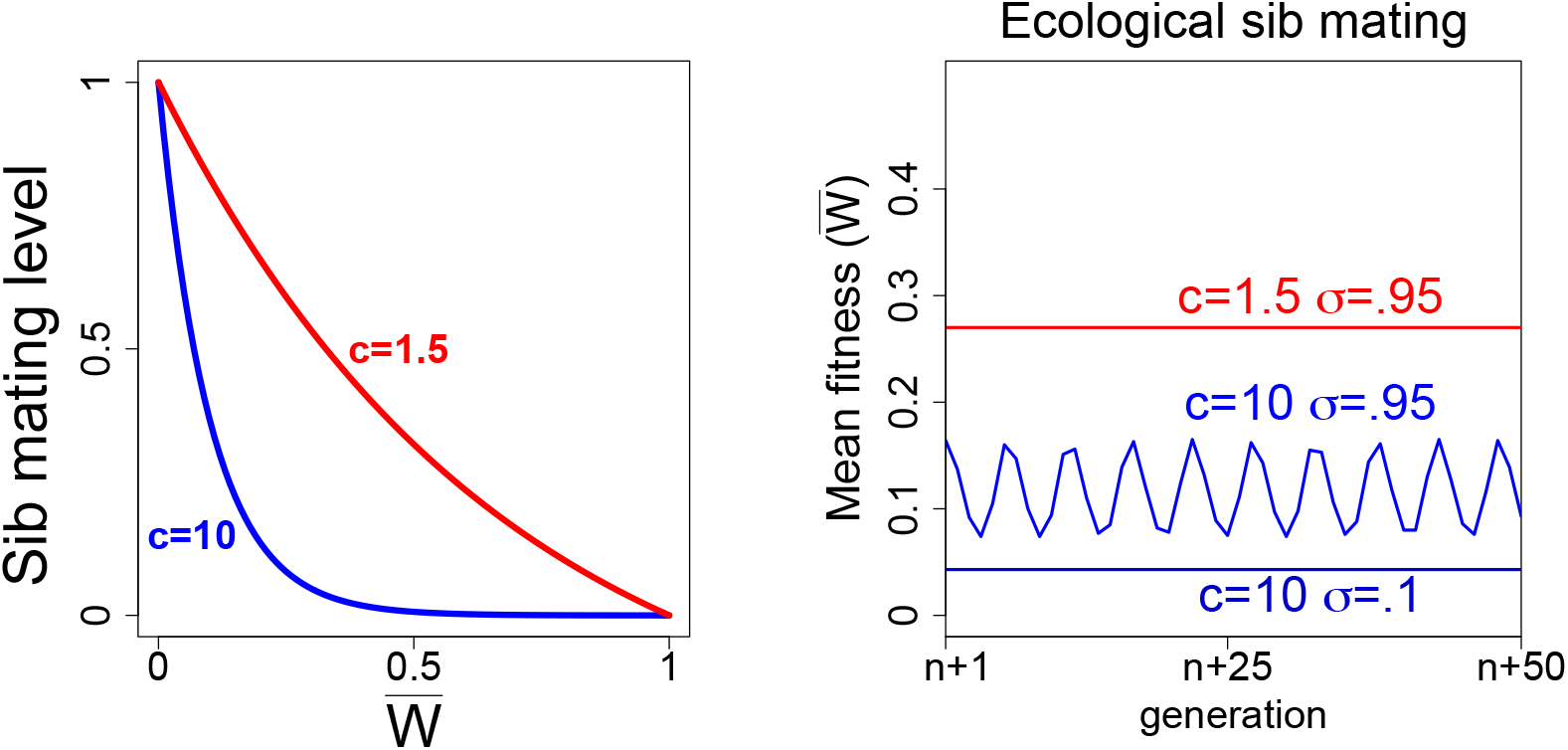
Environmental control of sib mating with recessive lethal drive. (Left). The sib mating function is shown for two different values of the shape parameter, *c*. (Right) Equilibrium outcomes for three different trials of the ecological sib mating model with a recessive lethal drive. The output shown spans 50 generations following following more than 1000 initial generations. Note that 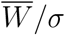 does even closely not approach 1 (this ratio must be inferred from the graph), but the ratio depends on *σ* and on the shape of the sib-mating function. Full male discounting was assumed (*K* = 0).

The evolutionary dynamics are intuitive: the drive allele spreads and depresses mean fitness, thereby increasing sib mating. The increase in sib mating affects spread of the drive allele, thereby limiting further drops in mean fitness or possibly increasing mean fitness. A balance may be reached – dynamic equilibrium – or oscillations may result (fig. 4). The shape of the sib mating function is critical both to the equilibrium as well as to the outcome of oscillations versus static equilibrium. Mean fitness depends heavily on the shape of the sib-mating function and again on *σ* – the fitness of inbred progeny.

### Y drive

#### Genetic control of sib mating

The model setup is similar in many ways to that of recessive lethal drive (Fig. 5). There are 4 types of sib mated families and 8 types of outcrossed families. The Y drive allele (Z chromosome) is present in four types of outcrossed families, and as those families produce only sons, there is no opportunity for sib mating. All sons carrying the Z allele join the outcrossing pool.

**Figure 5:**
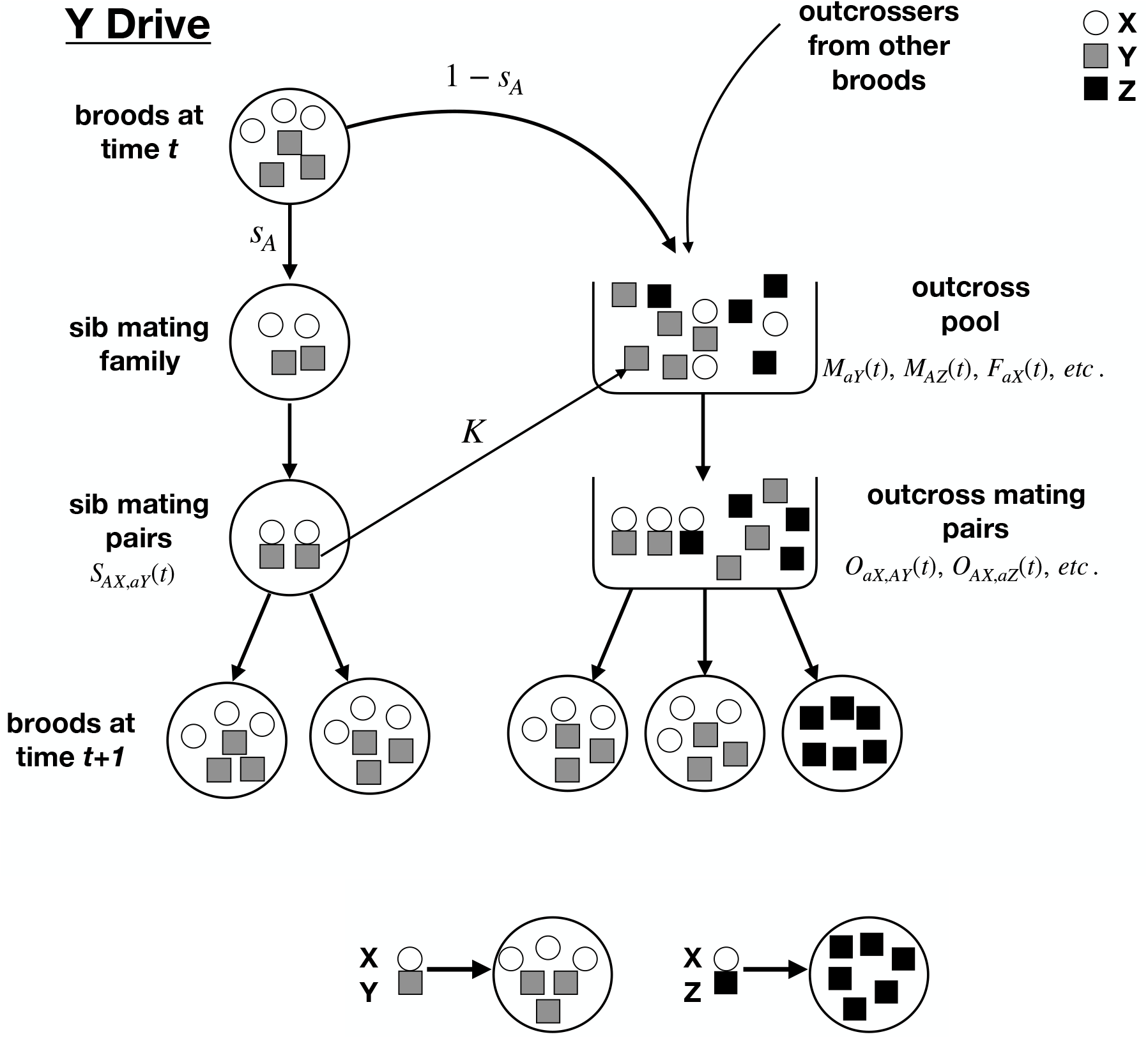
Schematic of the Y drive model with haploid individuals that enjoy a brief diploid phase for mating. Females are circles, males are squares; gray indicates a normal male (Y), black a male with the drive allele (Z). Top: Shown in the pedigree of large circles on the left is the life cycle of a brood that resulted from a mating of an *aX* female and an *aY* or *AY* male. The female sib mating allele determines the fraction of the brood (here *s*_*A*_) that is sib mated and the fraction 1 – *s*_*A*_ that go to the outcross pool along with progeny from other broods that are available to outcross. A fraction *K* of males that sib mate a sister also joins the outcross pool to possibly mate some more. Bottom: The mating of a female (X) and a Y male results in half female and half male offspring; the mating of a female and a Z male (containing gene drive) results in all Z male offspring – 100% drive efficiency. Each brood from an *XY* mating is assumed to carry an equal number of males and females, and any splitting of the brood retains these proportions. The effect of *σ* is not shown.

##### Equilibrium

As with recessive lethal drive, sib mating was found to evolve under Y drive, the details differing somewhat from the case of recessive lethal drive. For example, allele ‘A’ always evolved to fixation (allele ‘*a*’ was for strict outcrossing). Mean fitness relative to *σ* closely paralleled the sib mating level (fig. 6). (Mean fitness was calculated the same as for recessive lethal drive.)

**Figure 6:**
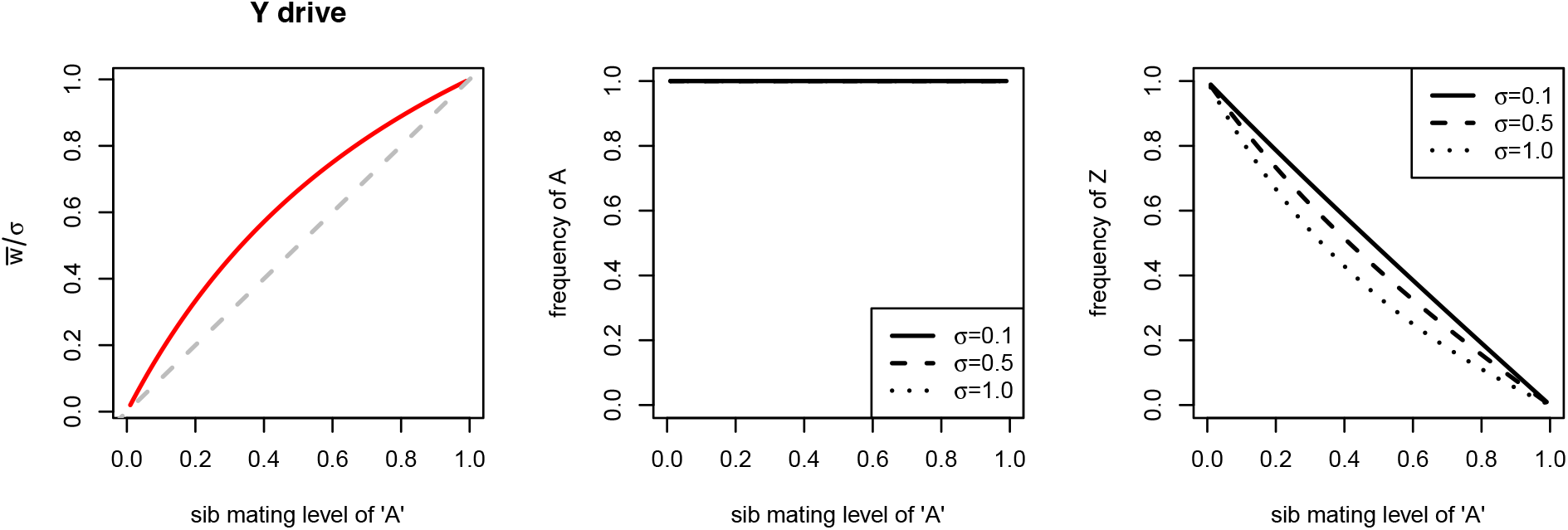
Genetic control of sib mating with Y (Z) chromosome drive: equilibrium. The horizontal axis in all panels is *s*_*A*_, the probability of sib-mating for allele ‘A’. All three panels are for the same trials, merely illustrating different properties. (Left) Relative mean fitness attained across different *s*_*A*_ values. 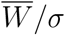 is mean fitness scaled by the fraction of maximum brood size attained by parents who were sibs. That this ratio was never observed to exceed one means that *σ* sets the upper limit on mean fitness, but in contrast to recessive lethal drive, here 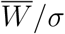 closely parallels *s*_*A*_ and never equals 1 except at *s*_*A*_ = 1. The dashed gray line is an isocline at which the y-value equals the x-value. (Middle) Equilibrium frequency of allele ‘A’. In contrast to recessive lethal drive, the ‘A’ allele always fixed. (Right) Equilibrium frequency of the drive allele, Z, closely follows 1 *- s*_*A*_. Trials assumed full male discounting (*K* = 0).

#### Ecological control of sib mating

Qualitatively similar results were obtained for ecological control of sib mating when assuming Y drive as when assuming recessive lethal drive. Oscillations appeared to require even more extreme deviations from linearity in the sib mating function under Y drive as under recessive lethal drive. Mean fitness remained well short of *σ*, even with a nearly linear function.

### Dynamics of all cases

Results described above are for long-term, equilibrium behavior. Short-term dynamics are of interest to understand how and how quickly equilibrium is attained. Fig. 7 shows representative dynamics from a single trial of each of the four classes of models. The most significant result is that genetic evolution of resistance experiences a large crash in mean fitness when the drive initially sweeps. This crash is due to the low initial frequency of the sib-mating ‘A’ allele, and the rebound in mean fitness is rapid. Nonetheless, the nadir in mean fitness is a vulnerability to extinction. Populations with the environmental inbreeding function do not experience this crash.

**Figure 7:**
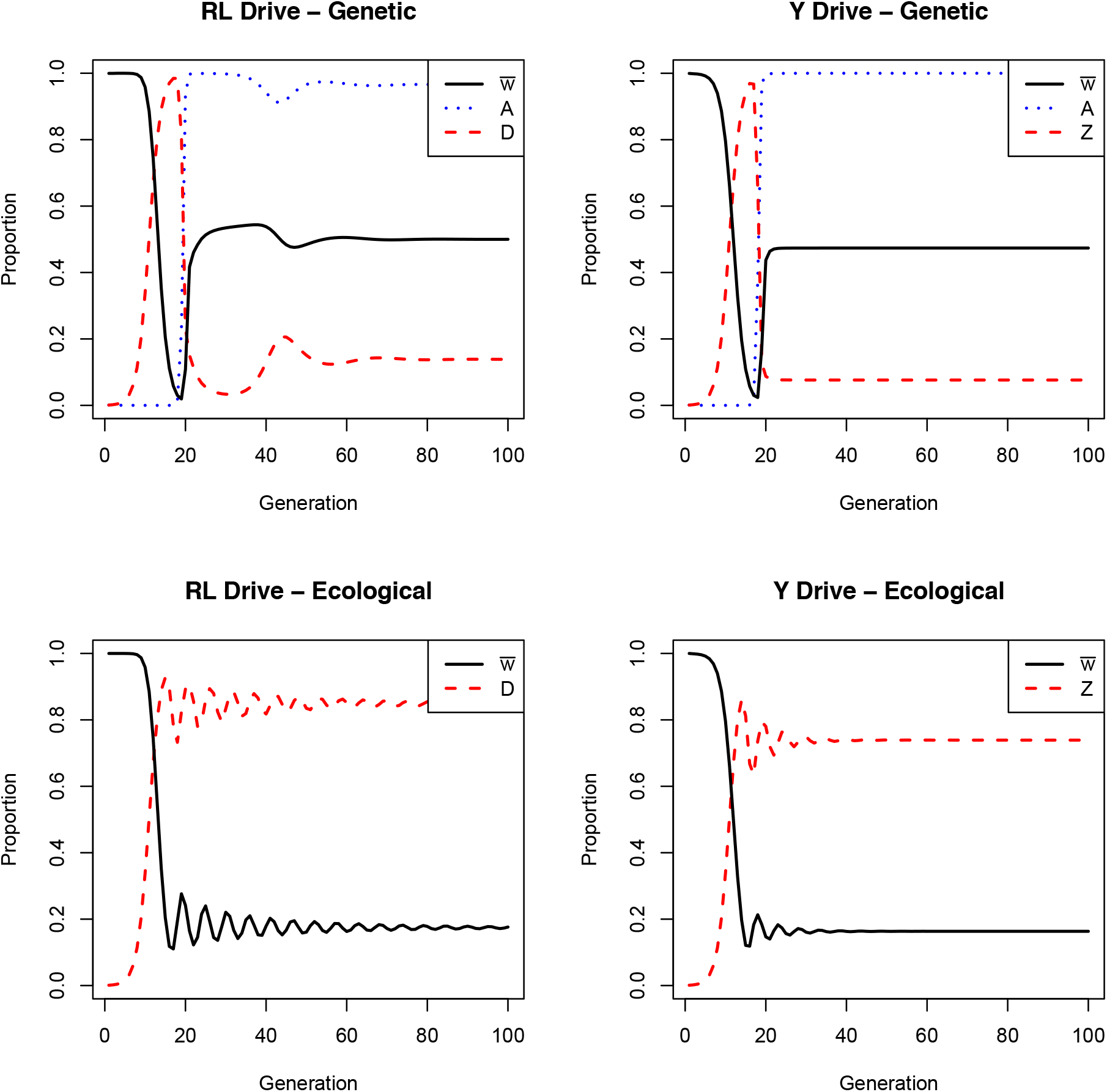
Dynamics of sib-mating rescue of extinction-causing gene drives: recessive-lethal (RL) drive and Y-chromosome drive. Black curves are mean fitness (bounded by 1.0), red are of the drive allele. Dotted curves in the top panels give the frequency of the sib-mating allele. Rescue by genetic control experiences an early dive in mean fitness, due to the allele for sib mating starting at a frequency of 0.001. Parameter values for genetic control of sib mating: *s*_*a*_ = 0, *s*_*A*_ = 0.9, *σ* = 0.5, *K* = 0. Parameter values for environmental control: *c* = 10, *σ* = 0.5, *K* = 0. Initial conditions: Recessive Lethal Drive – Genetic (*S*_*ad,Ad*_ = *S*_*Ad,ad*_ = 0.0005, *O*_*ad,aD*_ + *O*_*aD,ad*_ = 0.001, and *O*_*ad,ad*_ = 0.998); Recessive Lethal Drive – Environmental (*O*_*ad,aD*_+*O*_*aD,ad*_ = 0.001, *O*_*ad,ad*_ = 0.998); Y Drive – Genetic (*S*_*aX,aY*_ = 0.001, *O*_*aX,aZ*_ = 0.001, *O*_*aX,AY*_ = 0.998); Y Drive – Environmental (*O*_*aX,aZ*_ = 0.001, *O*_*aX,aY*_ = 0.999).

## 4 Discussion

The intentional engineering of gene drives has become so feasible that this intervention can be entertained for nearly any sexual plant or animal species. The two most basic possible uses of a gene drive system are to drag a genetic cargo through a population (with potentially little fitness consequence) or to suppress population reproduction, possibly to extinction. Extinction is the most profound and far-reaching of these applications, and it is also the most likely to select resistance.

Resistance evolution to block an intentional gene drive release will nearly always be undesirable, except perhaps in limiting the drive’s spread beyond the intended species. Even so, some types of resistance evolution will be far more undesirable than others, as some types of resistance will block future efforts that use the same or related technology.

Three classes of resistance evolution can be anticipated:

(R1) Resistance blocks the mechanism or action of the drive; with CRISPR, this resistance could involve changes in the nuclease target sequence, could block CRISPR expression, or interfere with the CRISPR RNA/protein complex.
(R2) Resistance may not interfere with the drive but merely compensate for its effects. Lyttle (1981) observed a change in *Drosophila* sex determination that tolerated the Y chromosome in both sexes and thus blocked the effect of a driving Y chromosome without blocking the drive. Burt (2003) noted that the effects of a recessive lethal drive could be abrogated by compensatory evolution in a different gene that assumed the function of the targeted gene.
(R3) As studied here, resistance may alter the mating structure of the population, protecting subsets of individuals from invasion by the driving element.

Type R1 underlies many mathematical models of resistance evolution, in that the resistant allele segregates opposite the drive allele (Burt 2003; Deredec, Burt, and Godfray 2008; Unckless, Clark, and Messer 2017). Type R1 resistance seemed like an inevitable outcome of lethal gene drive efforts using CRISPR (Champer et al. 2017; Drury et al. 2017), although the deployment of multiple guide RNAs to several targets at once was a possible bypass (Burt, 2003). However, a recent study identified a target sequence in mosquitoes that is both essential and apparently intolerant of change: caged populations of several hundred mosquitoes did not evolve resistance, instead going extinct (Kyrou et al. 2018). Mutations in the target sequence were observed at some life cycle stages, likely a consequence of imperfect repair of DNA lesions, but they were incompatible with fertility and thus did not evolve.

The potential for resistance – and types of resistance – will depend on the design of a drive system, as shown at least by Kyrou et al. (2018). The choice of a target sequence is obviously important. Some expression constructs may be more easily blocked than others. In addition, drives with cargo may be prone to lose the cargo, thereby introducing a secondary drive that competes with the original design but fails to provide the desired properties. Drives may even be engineered in various ways to prey on other drives, creating a type of arms race that will prevent any one from spreading throughout the population (Gantz and Bier, 2016).

The type of resistance that evolves has consequences beyond the immediate gene drive implementation. Specifically, there are different degrees to which resistance will be independent of the molecular mechanism of a gene drive. If resistance is a change in target sequence, that resistance will not affect drives that use other target sites. In contrast, resistance that is somewhat independent of the drive implementation may block future gene drive implementations that target other sites. Thus, resistance in the form of inbreeding and other changes in mating structure will block population-suppressing drives regardless of the technology. Inbreeding that is only partial may remain effective in limiting future population-suppressing drives but still allow the spread of harmless, cargo-carrying drives. The downside of any evolution of resistance that prevents extinction, even one in which mean fitness remains low, is that it provides a population nucleus in which further evolution of resistance may occur. Following evolution of inbreeding that even partially rescues, subsequent evolution could be of reduced inbreeding depression (Porcher and Lande, 2005; Charlesworth and Willis, 2009; Porcher and Lande, 2013) or could be other classes of resistance. Being non-essential, CRISPR may be especially prone to suppression in the long term. Rapid extinction may be the only hope for a persistent suppression.

The results here indicate that evolution of sib mating is a threat to an extinction-causing gene drive. Evolution of inbreeding is not assured, and its evolution is far more complicated than is evolution of a change in the target sequence. Even if inbreeding does evolve, the recovery of mean fitness is approximately limited by the magnitude of inbreeding depression. However, other forms of inbreeding are theoretically possible (e.g. Crow and Kimura, 1970), and some of these may be less affected by inbreeding depression than is sib mating.

### Anomalies in the theory

A previous theoretical analysis of the evolution of selfing in response to a lethal gene drive observed that selfing was favored only if the selfing allele enacted a sufficiently large degree of selfing (Bull, 2016). That was an encouraging result in suggesting that selfing could not evolve gradually. But that result was demonstrated for models of a recessive lethal drive in which drive was limited to one sex (and was not evaluated for models of drive in both sexes). None of the results here support that outcome.

Further numerical studies of those selfing models (conducted for the current study) suggest that the block to gradual evolution of selfing is due to a combination of recessive lethal drive, and (ii) incomplete drive (e.g., drive operating only in one sex, or the segregation distortion of the drive being less than 100%). An intuitive argument for that previous result and for our failure to observe it in the present recessive-lethal model analyses is as follows: when a drive distortion is complete, any mating in which one parent carries the drive allele produces a family in which every offspring carries the drive allele. The alleles for inbreeding in this family never return to the pool of genotypes that lack drive. But when drive is incomplete, a mating in which one parent carries the drive allele sometimes produces descendants that do not carry the drive allele, and they or their descendants can then return to the drive-free pool and influence the frequency of alleles for inbreeding. In such families, an allele that imposes a low level of inbreeding will return more progeny to the drive-free pool than does an allele that imposes a high level of inbreeding – because the drive allele is a recessive lethal. Thus, an incomplete drive partly selects against inbreeding by disproportionately returning low-inbreeding alleles back to the drive-free pool of genotypes.

We conjecture that similar arguments apply to some of the differences in evolutionary dynamics observed here between Y drive and recessive lethal drive (e.g., comparing figs 3 and 6). In particular, the mating between a parent carrying a recessive lethal drive allele and an allele for high inbreeding kills grandchildren from the inbreeding, whereas there is no such killing of grandchildren with Y drive. This asymmetry may explain why the sib-mating allele always fixes with Y drive but not always with recessive-lethal drive.

### Extinction

The models in this study are strictly of population genetics and do not account for population size. Yet the motivation for this work is to understand resistance that blocks extinction. Inferring extinction from mean fitness requires insight to ecology – fecundity, the nature of population regulation, Allele effects at low density, and so on. Predicting extinction will thus face many challenges beyond merely predicting resistance evolution. These difficulties are foreshadowed by a 1977 symposium on the evolution of resistance to the US implementation of the sterile insect technique against the screw worm (Richardson, 1979). The program had suffered a recent rebound in screw worm cases, and various forms of resistance evolution were entertained to anticipate a long term failure of the program. In the final analysis, the rebound was apparently due chiefly to factory evolution of the strains used for release; replacement of those strains in rearing facilities was adequate to restore program efficacy and led to ultimate eradication from North and Central America. Resistance evolution did not prevent program success despite the plausibility of various possible avenues of failure.

Yet even if predicting resistance evolution proves elusive, an understanding of how resistance might evolve makes it possible to take steps to identify the best candidate species for release and to ensure that failures of the first efforts do not thwart later efforts. Experience with the sterile insect technique led to the realization that some species characteristics were more prone to success than others (Klassen and Curtis, 2005; Itô and Yamamura, 2005). We can likewise suggest a few characteristics that should facilitate extinction by gene drive:

1. High inbreeding depression. If sib mating evolves, the fitness recovery is limited by inbreeding depression. Species with high inbreeding depression, and also for which inbreeding depression is slow to reverse on extended inbreeding, should face difficulty in escaping extinction through inbreeding evolution. Magnitudes of inbreeding depression are easily studied experimentally with any species that can be housed artificially, so species anticipated as targets of extinction-causing gene drives might be screened in advance to decide on the feasibility of inbreeding evolution.
2. Low fecundity. Declines in mean fitness have a greater impact in suppressing numbers of individuals when females produce few offspring than when they produce many. The nature of population regulation also enters into this effect.
3. Small population size or low density. Opportunities for resistance mutations should be fewer in small populations, and mating opportunities of any resistant individuals should also decline with population density. This principle was an important one behind success of the sterile insect technique.
4. Intrinsic outbreeding. Some life histories may be intrinsically disposed to outbreeding and thus face difficulty in evolving inbreeding.

Whether extinction-causing gene drives will commonly avoid resistance evolution remains to be seen. Nor is it clear that experience with the sterile insect technique will translate to gene drive extinctions: overwhelming a wild population with sterile individuals will have different consequences for the evolution of resistance than will suppression of population densities that creates a paucity of opportunities for mating. From this perspective, a Y chromosome drive may create a demography of extinction more similar to the sterile insect technique than does a recessive lethal drive. However, and as foreshadowed by the sterile insect technique, it is expected that some gene-drive extinction efforts will succeed and others will fail. We may at least hope that we can develop a sense for the difference and understand how to improve the chances of success.

## Acknowledgments

Mathematica 11.3.0.0 was used in some analyses. Supported by NIH GM 122079.

## Appendix

### Recessive lethal drive equations

Figure 2 provides a schematic to suggest some (but not all) model components.

#### Female haplotypes

*ad, Ad, aD, AD*; **Male haplotypes**: *ad, Ad, aD, AD*

(Any mating between a *D* male and a *D* female will be nonviable. All other matings produce offspring.)

Maximum **brood size** per mating: *b*

(Each outcrossed female produces a brood of *b* offspring, half of which are female. Each sib-mated female produces a brood of *σb* offspring, half of which are female.)

#### Mating frequencies for parents in generation *t*

When denoting mating pairs below, the first subscripted haplotype is for the female and the second is for the male. Quantities (appropriately subscripted) of the form *S*(*t*) and *O*(*t*) are fractions corresponding to (viable) mating pairs formed by the parents in generation *t*. These fractions add to 1. The offspring from these generation *t* matings have their future (generation *t* + 1) mating numbers encapsulated in (unnormalized) quantities of the form *S’*(*t*) and *O’*(*t*). The sum of all these “primed” quantities is the total, *T* (*t*), number of female offspring from generation *t* (the potential mothers of generation *t* + 1). Not all of these matings produce viable offspring. Dividing the *S’*(*t*) and *O’*(*t*) terms by *T* (*t*) produces the normalized quantities *S*(*t* + 1) and *O*(*t* + 1) of the next generation. The mean fitness in generation *t* is defined to be 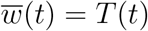.

Fractions of sib mating pairs:

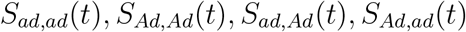

Fractions of outcrossed mating pairs:

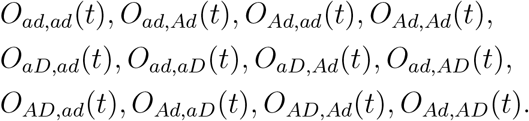

(For example, a fraction *S*_*Ad,ad*_(*t*) of the adult females in generation *t* have haplotype *Ad* and mate with an adult male (brother) with haplotype *ad*. These two first appeared as offspring in generation *t -* 1. Note that we do not list potential mating pairs with both parents carrying the D allele (e.g., *S*_*aD,aD*_(*t*) and *O*_*AD,aD*_(*t*)) since they produce no offspring.)

Numbers of outcrossed females: *F*_*ad*_(*t*), *F*_*Ad*_(*t*), *F*_*aD*_(*t*), *F*_*AD*_(*t*)

Fractions of outcrossed males: *M*_*ad*_(*t*), *M*_*Ad*_(*t*), *M*_*aD*_(*t*), *M*_*AD*_(*t*)

Note that we track actual numbers (densities) of female types and only fractions of male types. This is because each female is allowed to reproduce only once per generation but males can reproduce multiple times within a generation. Since offspring numbers are limited by the number of females (and not by the number of males), it is natural to track mean fitness in terms of females.

The offspring produced in generation *t* will be the adults of generation *t* + 1. Offspring and parents (as well as any unmated males) are assumed to coexist for a brief time in a given generation, but only offspring transition to the next generation (becoming the new adults); i.e., all individuals have a life span of one generation. To compute the mating quantities for the next generation, we first record the numbers of female **offspring** haplotypes and their *anticipated* mating types.

Future sib mated families (one per female; before normalization):

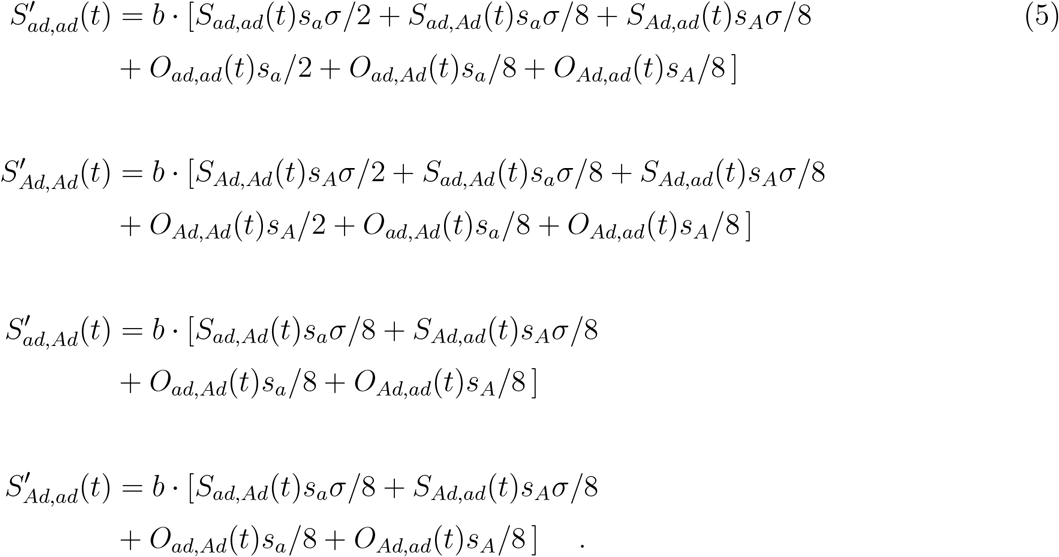

Future outcrossed families (one per female; before normalization): There are four equations of each of the following types

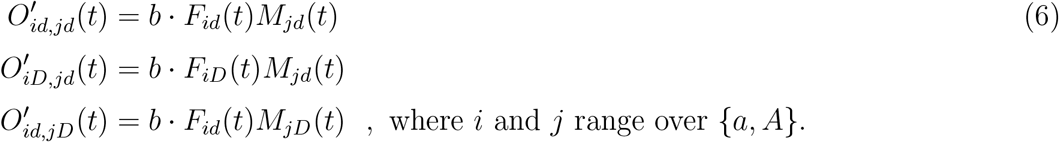

(Note that we could have included the nonviable mating pairs that can form, as long as we used brood size 0 *· b* instead of *b* (for viable outcrossed matings) or *σ* · *b* (for viable sib matings).)

Remark: The above equations are retrospective in the sense that quantifying the new mating types in equations (5) and (6) involves frequencies of mating types for females and males in the parent generation. To see where the different terms come from, it is instructive to think prospectively for a moment. For example, a fraction *S*_*Ad,ad*_(*t*) of adult females in generation *t* have haplotype *Ad* and sib mate with males (brothers) having haplotype *ad*. Since these are sib matings, they each produce a brood of size *σb*, half of which are female. Of these female offspring, all have allele *d* at the *d/D* locus (since both parents carried the *d* allele), and half have allele *a* at the *a/A* locus. Similar fractions apply for the male offspring. Of the *σb/*2 female offspring from each such mating, a fraction *s*_*A*_ will mate with a brother and a fraction 1 *- s*_*A*_ will outcross. (The mother’s haplotype at the *a/A* locus determines the fraction of her offspring that sib mate.) So, for example, each of the featured matings will produce *σb/*2 *× s*_*A*_ *×* (1*/*2) *×* (1*/*2) = *σbs*_*A*_*/*8 future matings to 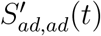.

The total

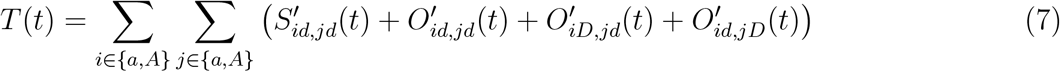

serves two purposes. Since parental mating pair frequencies are normalized, *T* (*t*) is the mean number of female offspring per female adult in generation *t*. This is **mean fitness** in generation *t*:

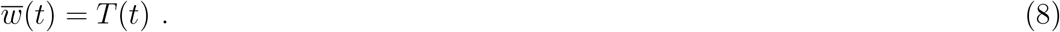

*T* (*t*) also serves as the normalizing quantity used to turn the raw counts of (anticipated) matings by generation *t* offspring into the normalized family frequencies at time *t* + 1.

#### Updating quantities for generation *t* + 1

Normalized family frequencies at time *t* + 1: There are four equations of each of the following types

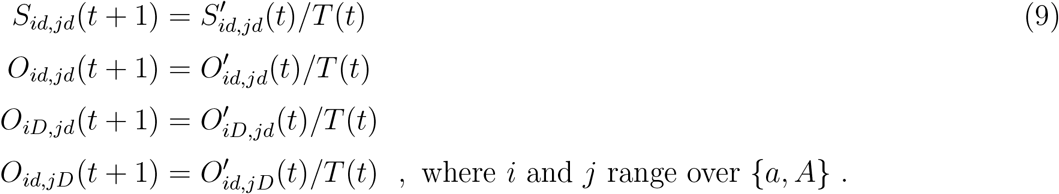

Notice that the maximum brood size, *b*, cancels in the above ratios. It does appear in mean fitness, though our plots of mean fitness were generated with *b* = 1; i.e., plots of mean fitness are normalized by brood size. In fact, the effect of brood size is transient in this model since we normalize down to mating *frequencies* each generation. Thus our model does not include population demography.

Outcrossed females at time *t* + 1 (never normalized):

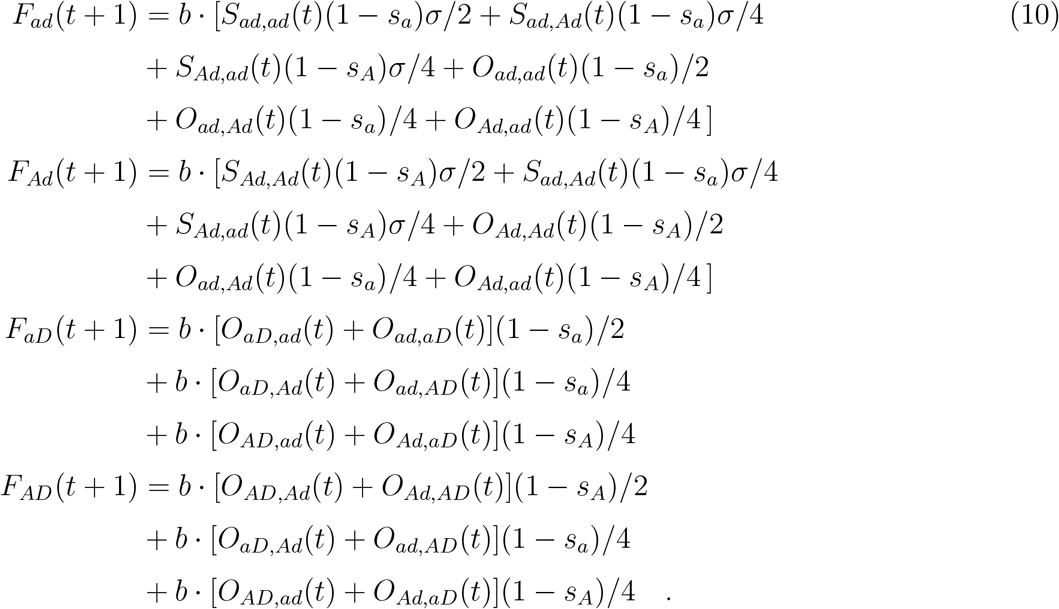

Outcrossed males (before normalization):

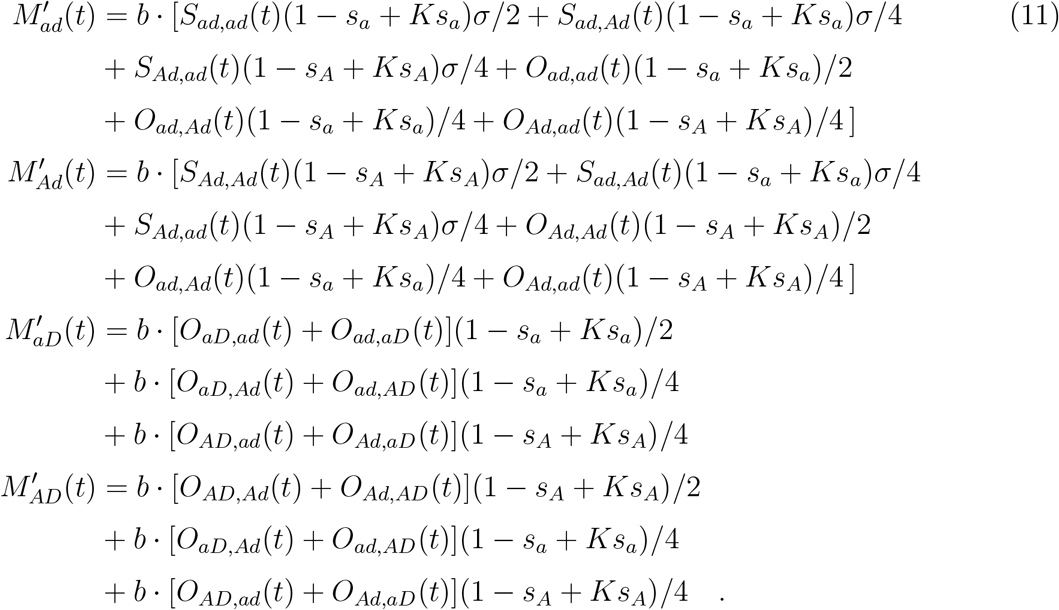

Normalized outcrossed male frequencies at time *t* + 1:

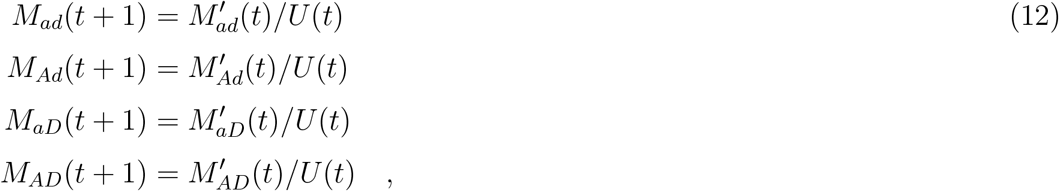

where 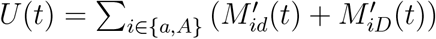.

### Y drive equations

Figure 5 provides a schematic to suggest some (but not all) model components.

Female haplotypes: *aX, AX*; Male haplotypes: *aY, AY, aZ, AZ*

(Z is the Y drive allele, carried only in males. Any mating involving a Z male results in all Z male offspring; no daughters. All matings involving a Y male results in a brood of half daughters and half Y sons. There can be no sib mating involving Z males since they have no sisters.)

Brood size is as in the recessive lethal model.

#### Mating frequencies for parents in generation *t*

When denoting mating pairs below, the first subscripted haplotype is for the female and the second is for the male. Other notations parallel those in the recessive lethal drive model description.

Fractions of sib mating pairs:

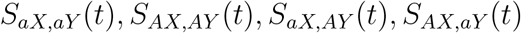

Fractions of outcrossed mating pairs:

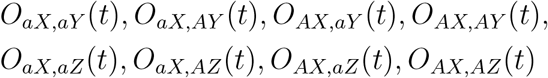

Numbers of outcrossed females: *F*_*aX*_(*t*), *F*_*AX*_(*t*)

Fractions of outcrossed males: *M*_*aY*_ (*t*), *M*_*AY*_ (*t*), *M*_*aZ*_(*t*), *M*_*AZ*_(*t*)

To compute the mating quantities for the next generation, we first record the numbers of female **offspring** haplotypes and their *anticipated* mating types.

Future sib mated families (one per female; before normalization):

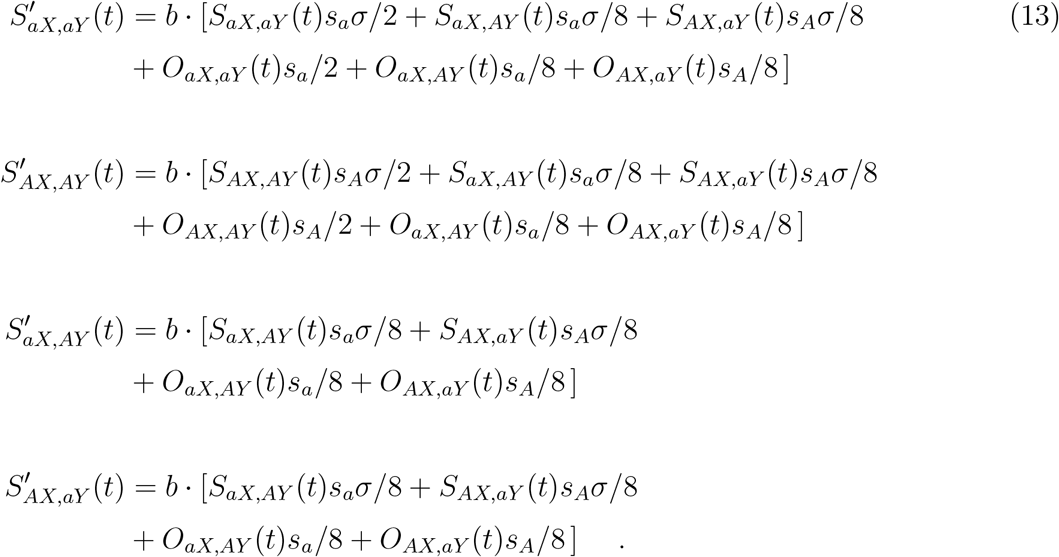

Outcrossed families (one per female; before normalization): There are four equations of each of the following types

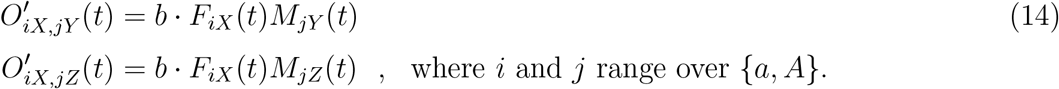

The total

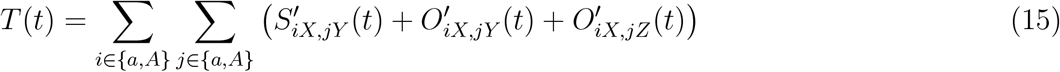

plays a role similar to the analogous total for the recessive lethal drive. **Mean fitness**in generation *t* is

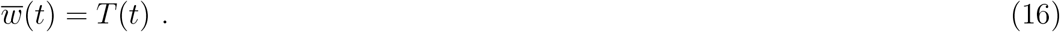

*T* (*t*) also serves as the normalizing quantity used to turn the raw counts of (anticipated) matings by generation *t* offspring into the normalized family frequencies at time *t* + 1.

### Updating quantities for generation *t* + 1

Normalized family frequencies at time *t* + 1: There are four equations of each of the following types

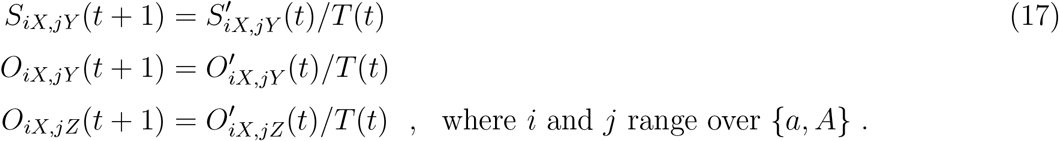

Outcrossed females at time *t* + 1 (never normalized):

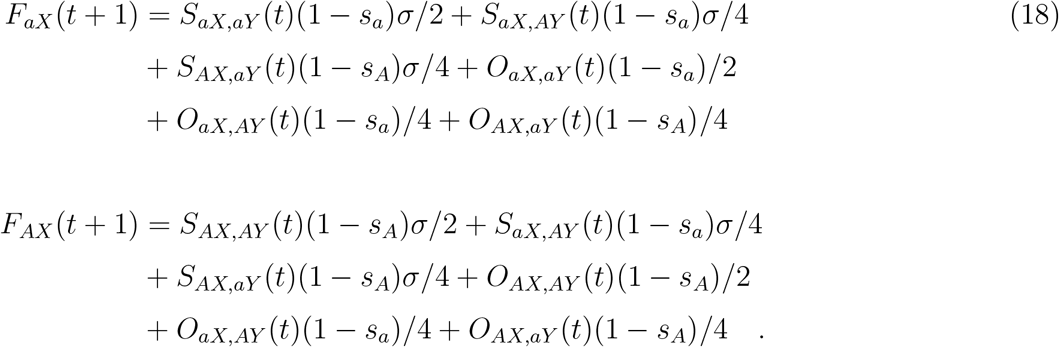

Outcrossed males (before normalization):

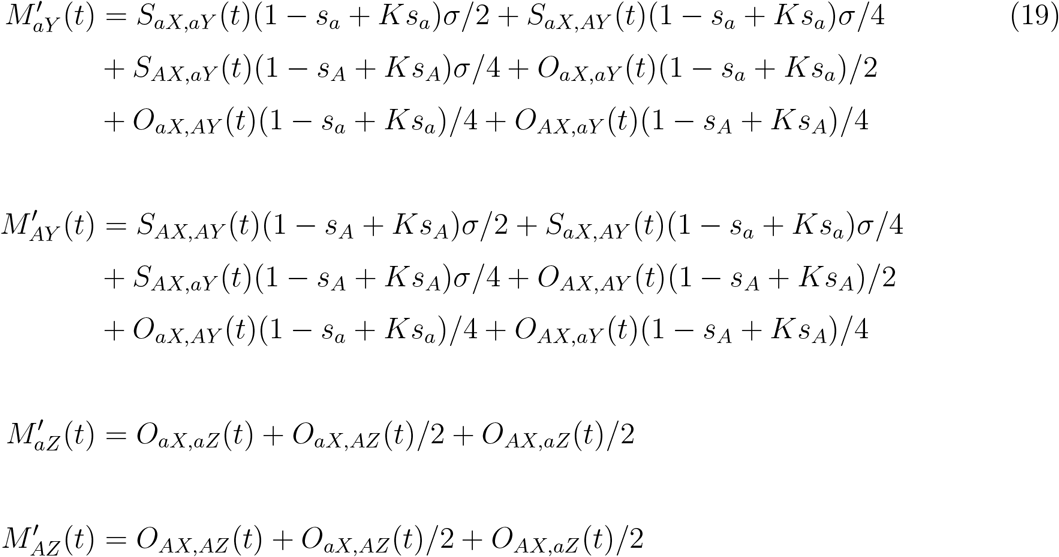

Normalized outcrossed male frequencies at time *t* + 1:

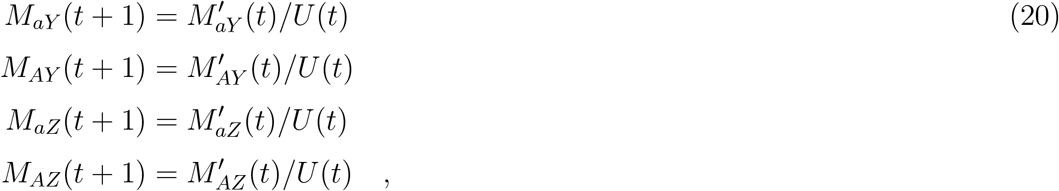

where 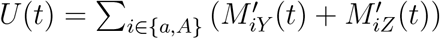.

#### Recessive lethal drive and Y drive with ecologically determined sib mating rates

In these models, there is only one sib mating frequency *s*_*a*_ = *s*_*A*_ and it is determined by mean fitness according to equation (4). Aside from this change in the dynamic updating of sib mating frequency, the equations for recessive lethal drive and Y drive are as before.

